## Supplemental figures and tables for "Divergent neural circuits for proprioceptive and exteroceptive sensing of the *Drosophila* leg"

### Supplementary Materials

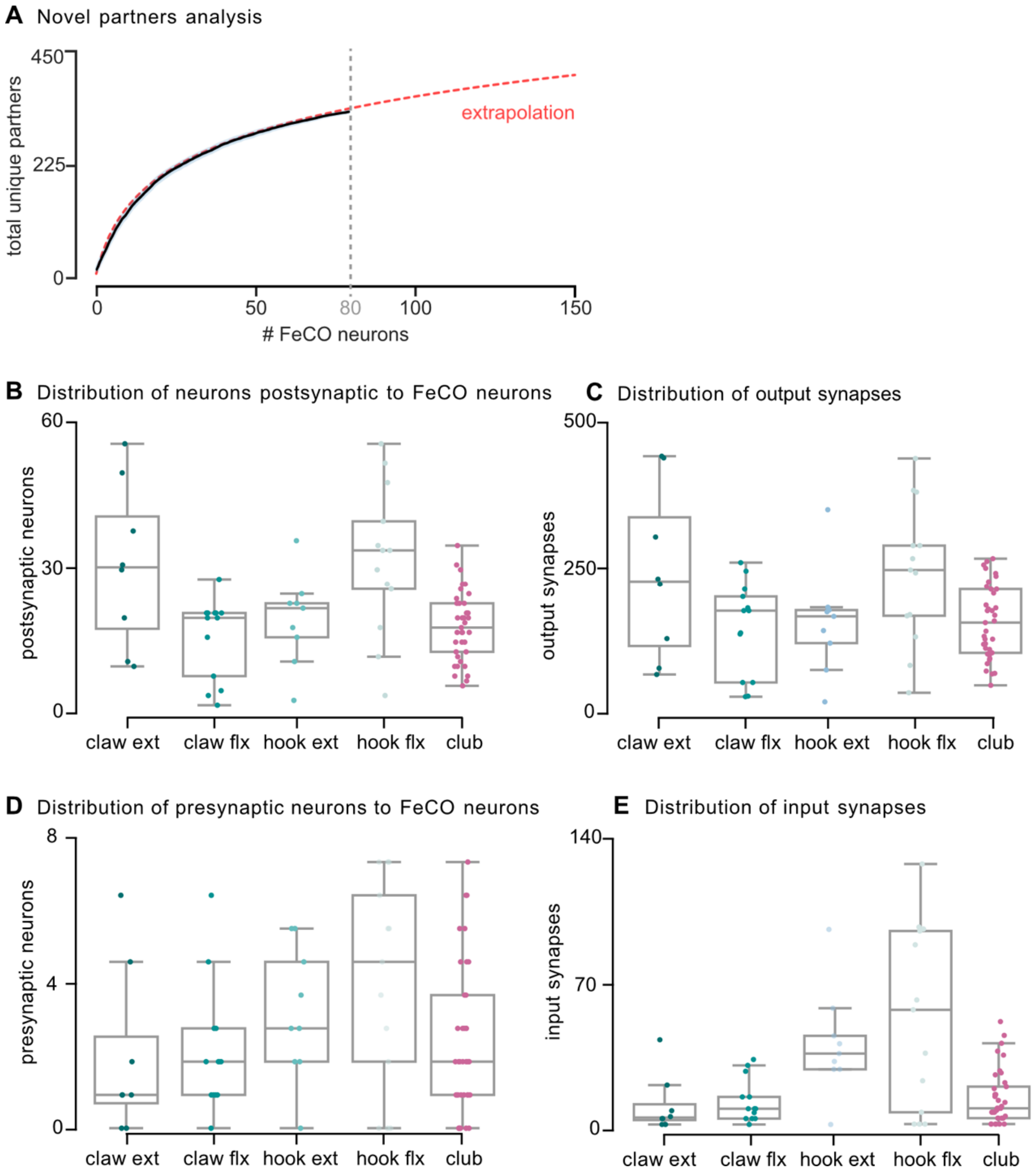


**Figure S1. Novel partners analysis suggests we have reconstructed a meaningful fraction of the FeCO axons from the front left leg.** (A) Plot shows the average number of new postsynaptic partners added per each FeCO sensory neuron we reconstructed. We randomly sampled the FeCO neurons one at a time (without replacement) in a cumulative fashion, and calculated how many novel postsynaptic partners were connected to each additional FeCO neuron. We resampled fifty times. Mean (black solid line) and standard error (shaded region surrounding line) are plotted. We then extrapolated the data (red dashed line) to estimate the remaining postsynaptic partners that have not yet been proofread. (B) Distribution of the number of unique neurons that are postsynaptic to each FeCO neuron subtype. The postsynaptic neurons include both proofread neurons and fragments that receive at least 4 synapses from a FeCO neuron. (C) Distribution of the number of synapses between each FeCO neuron and an individual postsynaptic partner. (D) Distribution of the number of unique neurons that are presynaptic to each FeCO neuron. The presynaptic neurons include both proofread neurons and fragments that make at least 3 synapse onto a FeCO neuron. (E) Distribution of the number of synapses between each FeCO neuron and an individual presynaptic partner.

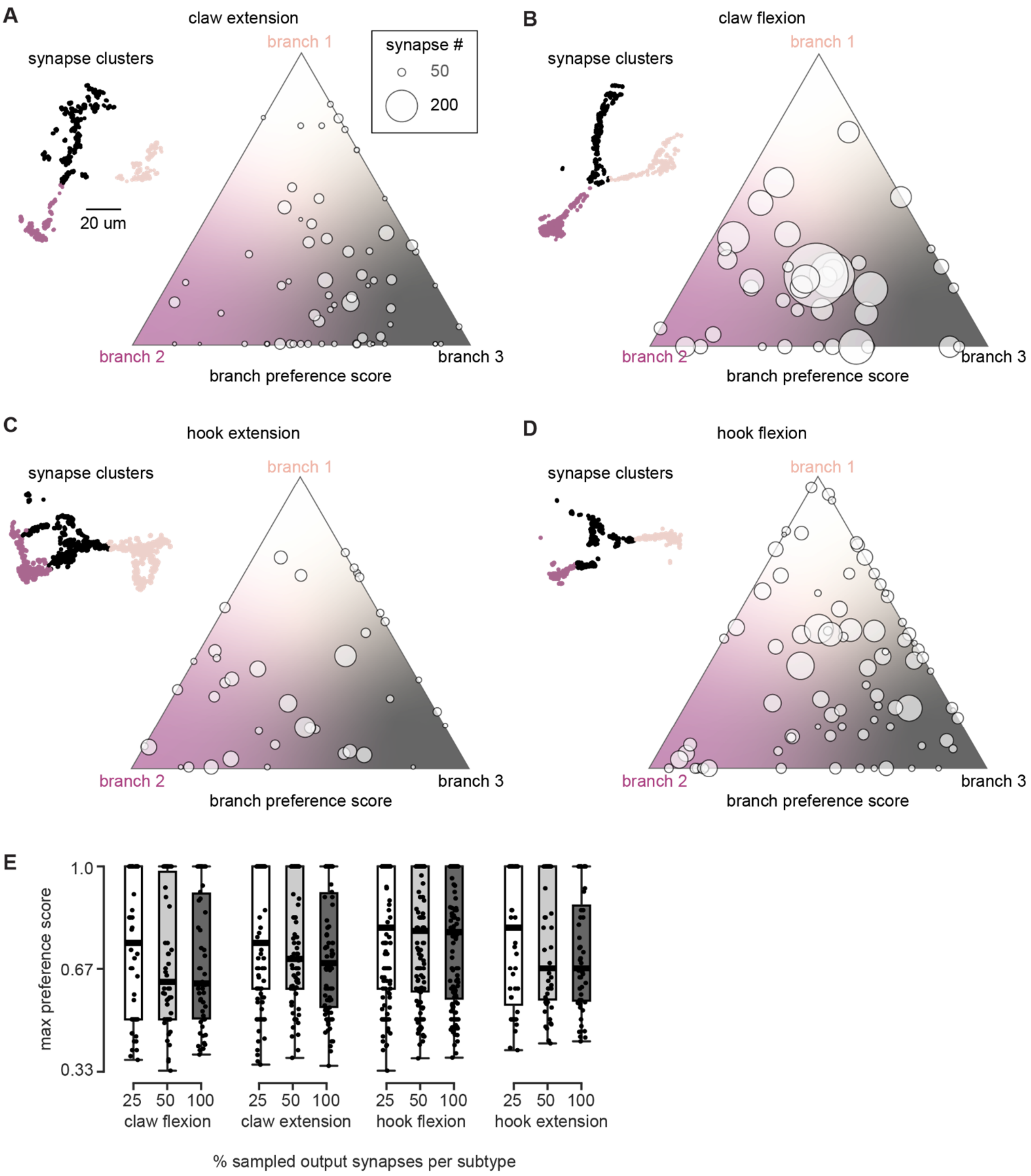


**Figure S2. The majority of postsynaptic neurons receive synaptic input from multiple branches of a given FeCO subtype.** A-D) We clustered the output synapses of each FeCO subtype based on their Euclidean distance (clusters shown on left). We then determined, for each postsynaptic partner, what fraction of its synaptic input came from which of these three clusters (branch preference score, see methods). Preference scores of each postsynaptic partner for each branch were then plotted on ternary plots (right). Size of the circle reflects the total number of input synapses the neuron receives from that FeCO subtype. E) We randomly subsampled either 25%, 50%, or 100% of an FeCO subtype's output synapses. We then calculated the branch preference scores of all postsynaptic partners. We plotted the max preference score per postsynaptic partner per FeCO subtype. Thick black line denotes the mean. No preference would be equivalent to a score of 0.33.

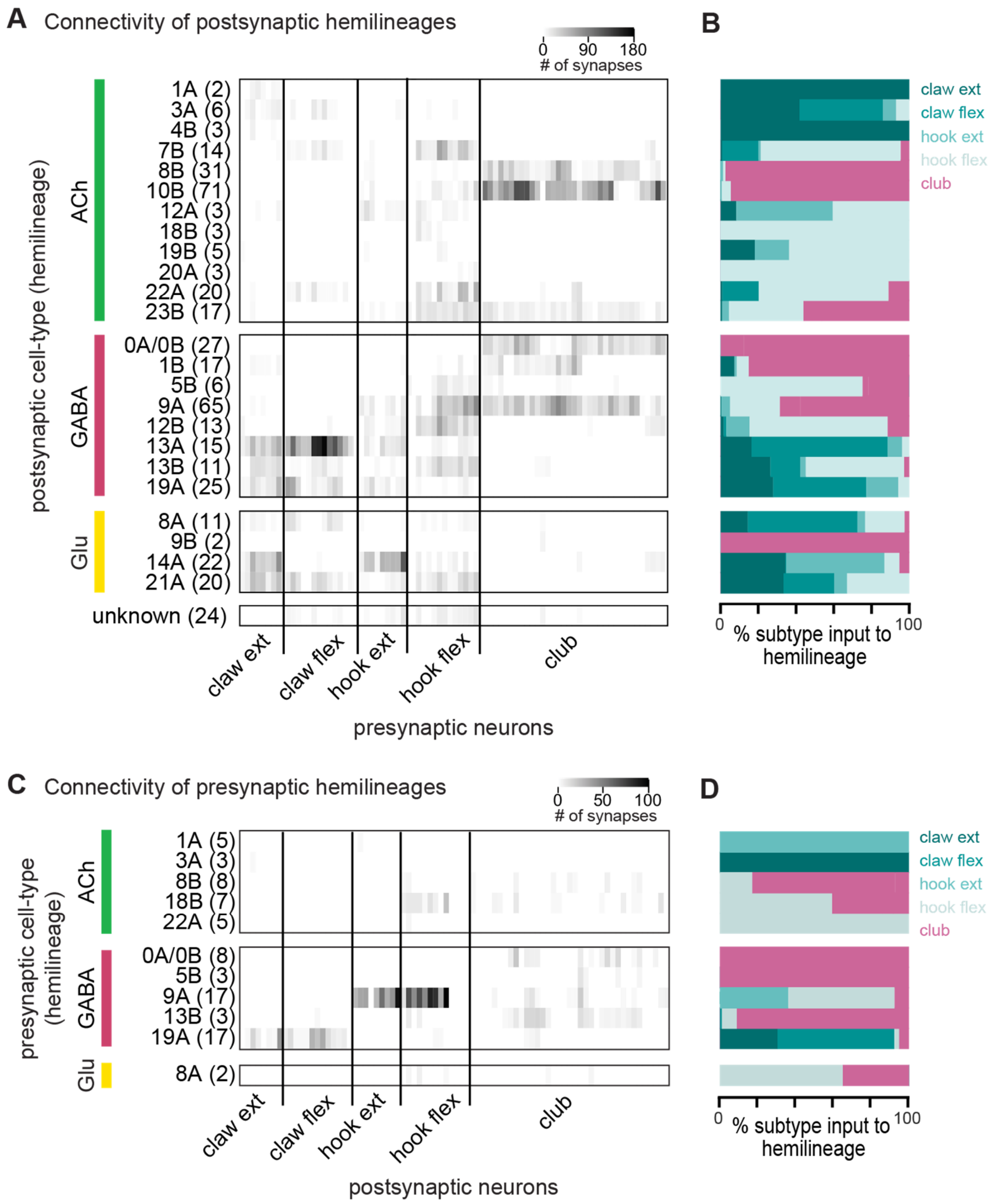


**Figure S3. Subtypes of FeCO axons preferentially synapse with VNC neurons from specific developmental lineages.** (A) The total number of synapses made by each FeCO neuron (columns) onto VNC neurons of each hemilineage (rows). Hemilineage was identified based on morphological characteristics of VNC neurons (see methods). Only local, ascending, and intersegmental postsynaptic partners are included in this analysis. VNC neurons of a given hemilineage are grouped together, with the number of neurons in each hemilineage indicated in parentheses. Hemilineages are grouped according to their primary neurotransmitter (ACh: acetylcholine, GABA: Gamma-aminobutyric acid, Glu: glutamate). (B) Percent of total FeCO input to a hemilineage made by each FeCO subtype. (C) The total number of synapses onto each FeCO neuron (columns) by presynaptic VNC neurons of each hemilineage (rows). (D) Percent of total inputs by a hemilineage onto each FeCO subtype.

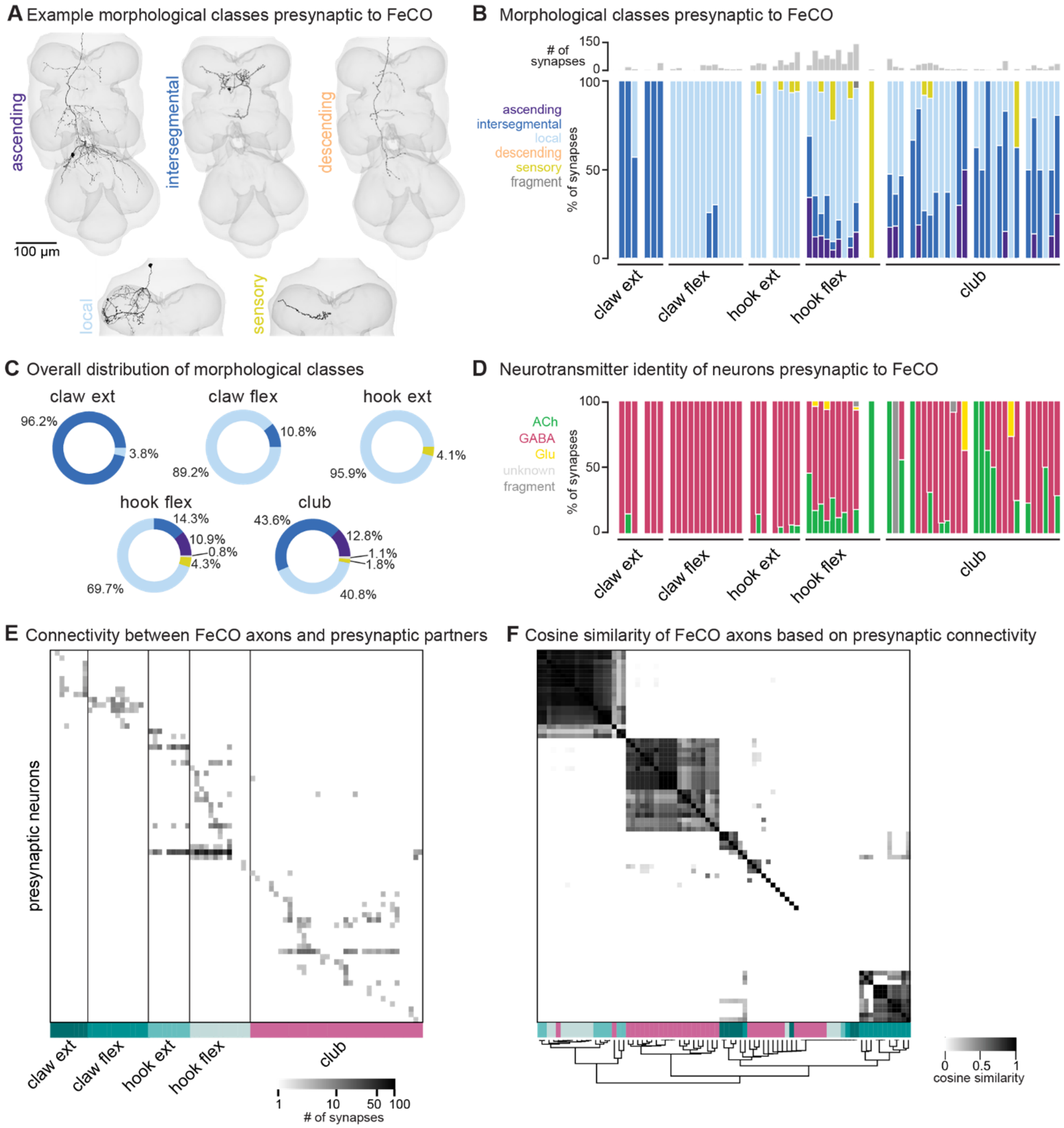


**Figure S4. FeCO neurons exhibit subtype-specific presynaptic connectivity.** (A) We reconstructed all VNC neurons presynaptic to FeCO axons from the front left leg (T1L) and classified them into morphological classes: ascending, descending, intersegmental, local, and sensory. Example provided from each class. (B) Percent of synapses received by each FeCO axon from VNC neurons of each morphological class. Top bar plot shows the total number of input synapses received by each FeCO axon. (C) Per FeCO subtype, the total fraction of input synapses received from each morphological class. (D) Proportion of total synapses made onto each FeCO neuron by presynaptic cholinergic (green), glutamatergic (yellow), GABAergic (pink), and unidentified (light gray) hemilineages. (E) Connectivity matrix between FeCO axons and presynaptic VNC neurons. The shading of each tick indicates the number of synapses from each presynaptic VNC neuron (row) onto each FeCO axon (column). Colored bars along the bottom indicate the postsynaptic FeCO subtype for that column. FeCO axons are organized by morphological subtype and then by their cosine similarity scores. VNC neurons are organized by their cosine similarity scores. (F) Clustered pairwise cosine similarity matrices of all FeCO axons based on their presynaptic connectivity. FeCO neurons with similar presynaptic connectivity patterns cluster together, forming connectivity clusters.

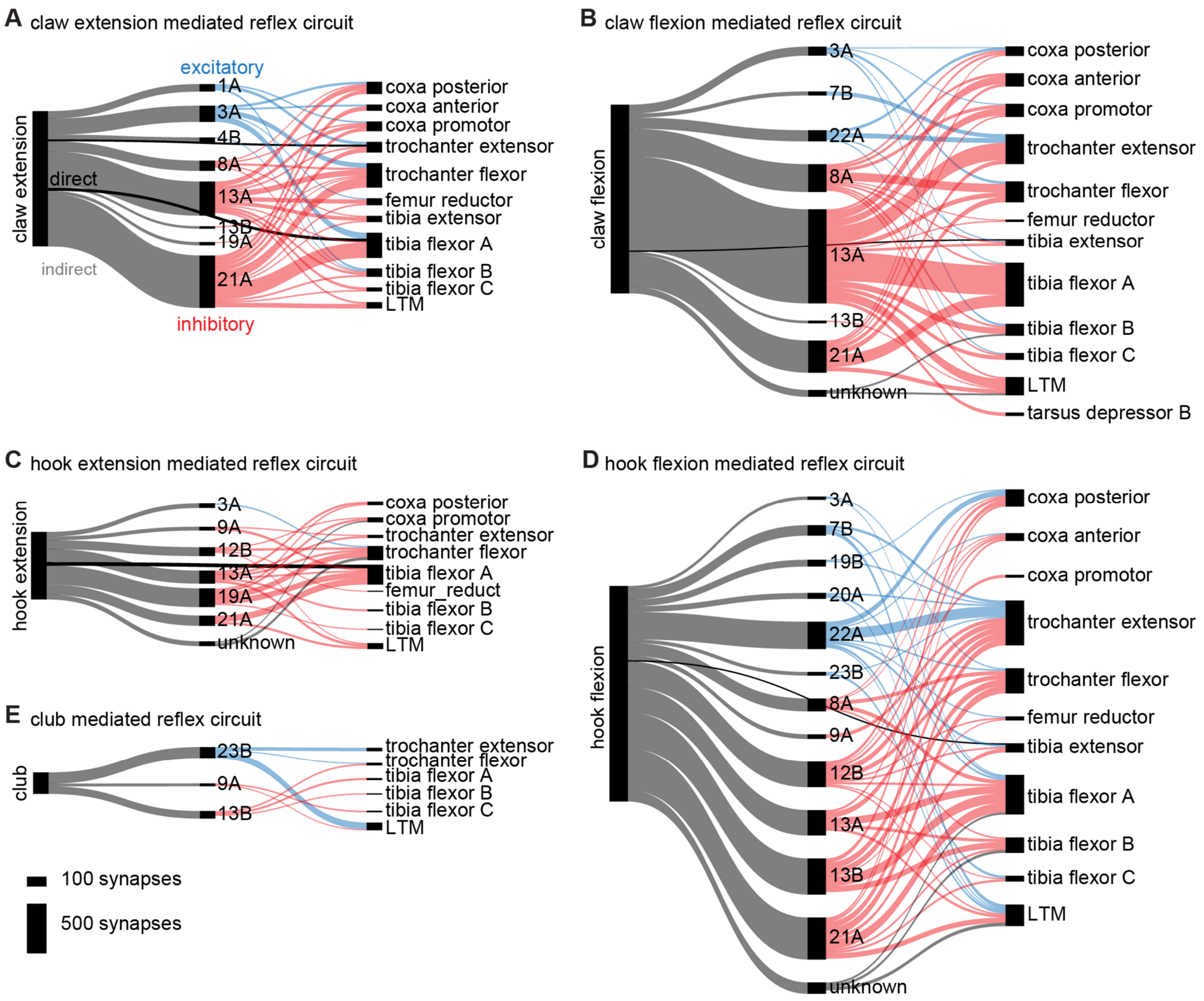


**Figure S5. Hook and claw neurons connect directly and indirectly to leg motor neurons.** FeCO neurons synapse onto leg motor neurons (black) and onto premotor neurons (gray). Leg motor neurons are grouped according to motor modules (Lesser et al., 2024) and premotor neurons are grouped by their developmental hemilineage (see methods). Premotor neurons were then identified as making either excitatory (blue), inhibitory (red) or unknown (gray) synapses onto leg motor neurons. Only premotor and motor neurons that received at least 10 synapses from FeCO axons and only premotor neurons that supply at least 10 synapses to motor neurons are displayed.

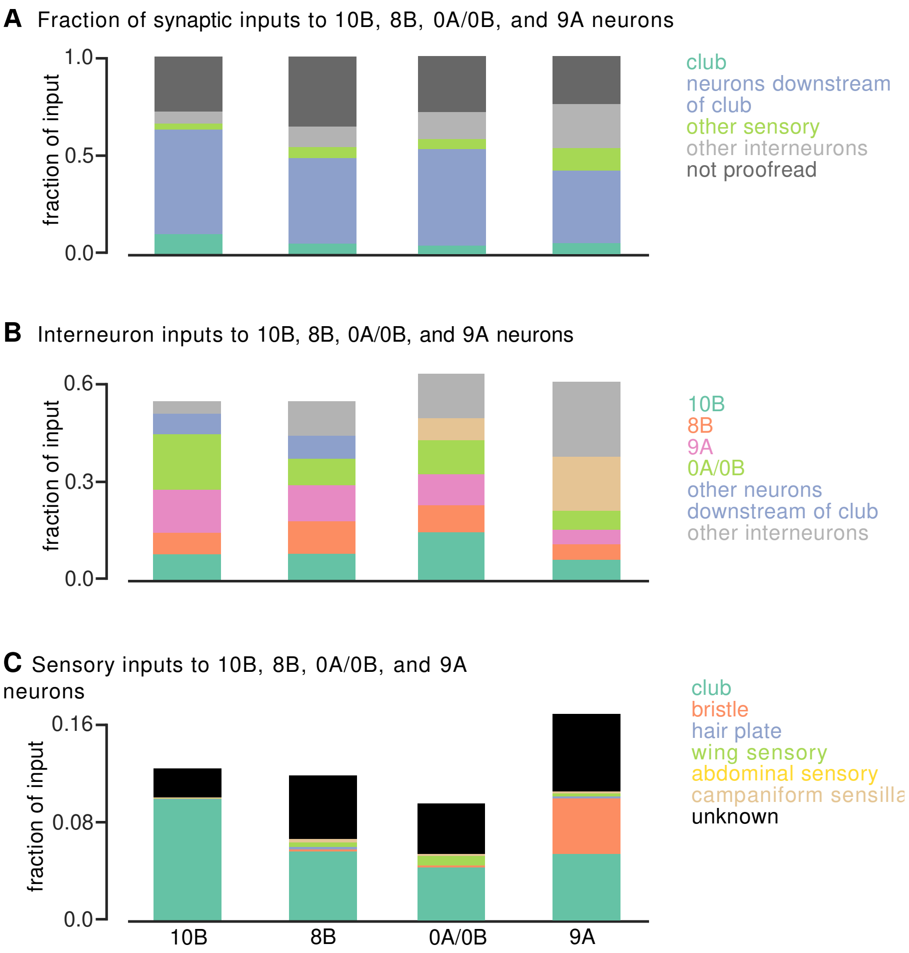


**Figure S6**. 8B and 10B neurons receive the majority of inputs from club circuits. (A) Fraction of total 10B, 8B, 0A/0B, and 9A input from club neurons, neurons downstream of club neurons (second-order and third-order interneurons), other sensory neurons, other interneurons, or neuron fragments. (B) Fraction of total input to 10B, 8B, 0A/0B, and 9A interneurons made by hemilineages that are postsynaptic to club neurons (10B, 8B, 9A, 0A/0B, other neurons downstream of club neurons, or other interneurons. (C) Fraction of total input to 10B, 8B, 0A/0B, and 9A interneurons made by sensory neurons.

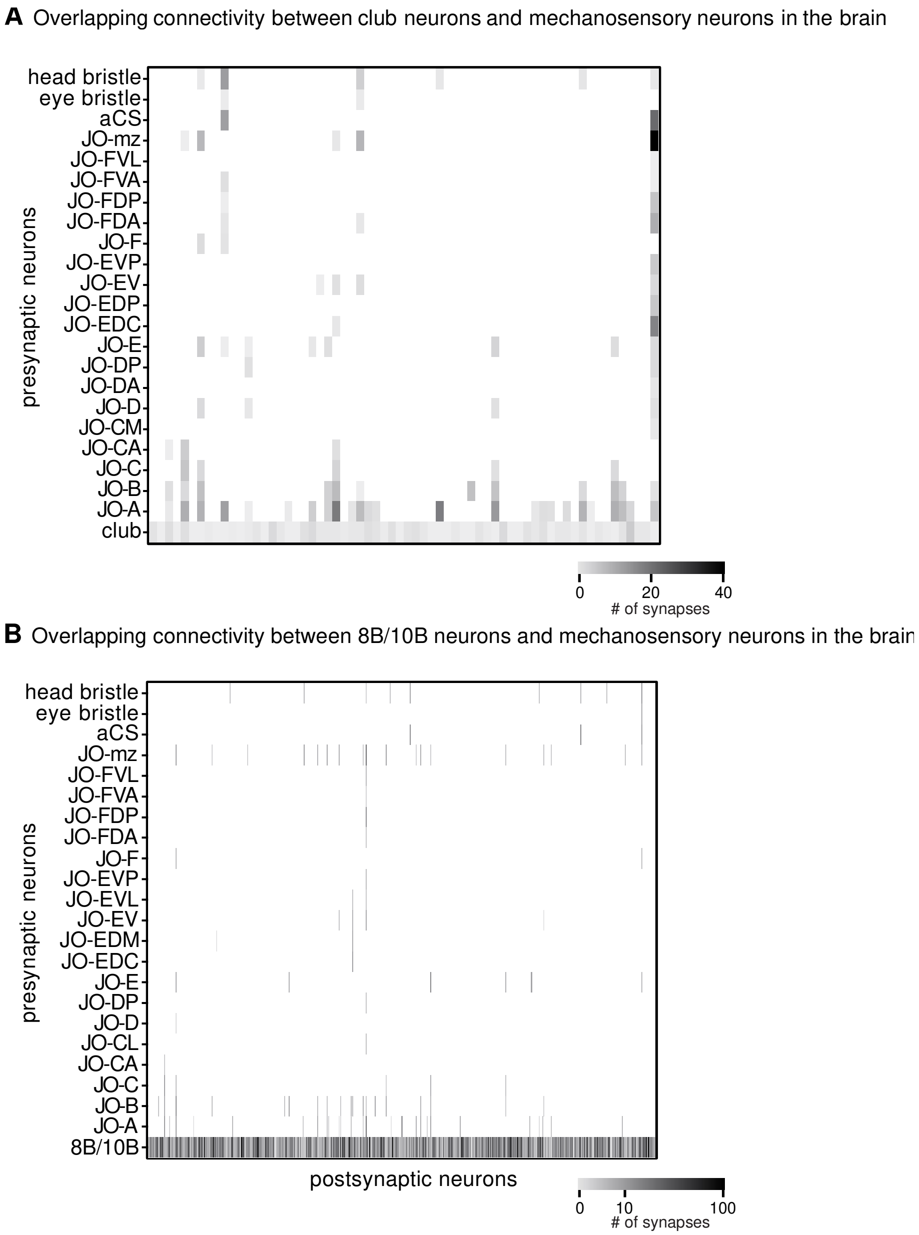


**Figure S7.** Ascending club neurons and 8B/10B neurons converge with mechanosensory neurons in the brain. (A) The total number of synapses made by club neurons and mechanosensory neurons (rows) onto postsynaptic neurons (columns) in the brain. (B) The total number of synapses made by 8B/10B neurons and mechanosensory neurons (rows) onto postsynaptic neurons (columns) in the brain.

**Supplemental Table 1.** Neuroglancer links to FeCO axons by subtype

| | |
|---|---|
| Club | https://neuromancer-seung-import.appspot.com/?json_url=https://raw.githubusercontent.com/sagrawal/Lee_2024/main/jsons/clubs.json |
| Hook extension | https://neuromancer-seung-import.appspot.com/?json_url=https://raw.githubusercontent.com/sagrawal/Lee_2024/main/jsons/hookE.json |
| Hook flexion | https://neuromancer-seung-import.appspot.com/?json_url=https://raw.githubusercontent.com/sagrawal/Lee_2024/main/jsons/hookF.json |
| Claw extension | https://neuromancer-seung-import.appspot.com/?json_url=https://raw.githubusercontent.com/sagrawal/Lee_2024/main/jsons/clawE.json |
| Claw flexion | https://neuromancer-seung-import.appspot.com/?json_url=https://raw.githubusercontent.com/sagrawal/Lee_2024/main/jsons/clawF.json |

**Supplemental Table 2.** Neuroglancer links to VNC neurons by hemilineage

| | |
|---|---|
| All presynaptic partners | https://neuromancer-seung-import.appspot.com/?json_url=https://raw.githubusercontent.com/sagrawal/Lee_2024/main/jsons/all_upstream.json |
| All postsynaptic partners | https://neuromancer-seung-import.appspot.com/?json_url=https://raw.githubusercontent.com/sagrawal/Lee_2024/main/jsons/all_downstream.json |
| Postsynaptic 0A/0B | https://neuromancer-seung-import.appspot.com/?json_url=https://raw.githubusercontent.com/sagrawal/Lee_2024/main/jsons/DS_0a0b.json |
| Postsynaptic 1A | https://neuromancer-seung-import.appspot.com/?json_url=https://raw.githubusercontent.com/sagrawal/Lee_2024/main/jsons/DS_1A.json |
| Postsynaptic 1B | https://neuromancer-seung-import.appspot.com/?json_url=https://raw.githubusercontent.com/sagrawal/Lee_2024/main/jsons/DS_1B.json |
| Postsynaptic 3A | https://neuromancer-seung-import.appspot.com/?json_url=https://raw.githubusercontent.com/sagrawal/Lee_2024/main/jsons/DS_3A.json |
| Postsynaptic 4B | https://neuromancer-seung-import.appspot.com/?json_url=https://raw.githubusercontent.com/sagrawal/Lee_2024/main/jsons/DS_4B.json |
| Postsynaptic 5B | https://neuromancer-seung-import.appspot.com/?json_url=https://raw.githubusercontent.com/sagrawal/Lee_2024/main/jsons/DS_5B.json |
| Postsynaptic 7B | https://neuromancer-seung-import.appspot.com/?json_url=https://raw.githubusercontent.com/sagrawal/Lee_2024/main/jsons/DS_7B.json |
| Postsynaptic 8A | https://neuromancer-seung-import.appspot.com/?json_url=https://raw.githubusercontent.com/sagrawal/Lee_2024/main/jsons/DS_8A.json |
| Postsynaptic 8B | https://neuromancer-seung-import.appspot.com/?json_url=https://raw.githubusercontent.com/sagrawal/Lee_2024/main/jsons/DS_8B.json |
| Postsynaptic 9A | https://neuromancer-seung-import.appspot.com/?json_url=https://raw.githubusercontent.com/sagrawal/Lee_2024/main/jsons/DS_9A.json |
| Postsynaptic 9B | https://neuromancer-seung-import.appspot.com/?json_url=https://raw.githubusercontent.com/sagrawal/Lee_2024/main/jsons/DS_9B.json |

| | |
|---|---|
| Postsynaptic 10B | https://neuromancer-seung-import.appspot.com/?json_url=https://raw.githubusercontent.com/sagrawal/Lee_2024/main/jsons/DS_10B.json |
| Postsynaptic 12A | https://neuromancer-seung-import.appspot.com/?json_url=https://raw.githubusercontent.com/sagrawal/Lee_2024/main/jsons/DS_12A.json |
| Postsynaptic 12B | https://neuromancer-seung-import.appspot.com/?json_url=https://raw.githubusercontent.com/sagrawal/Lee_2024/main/jsons/DS_12B.json |
| Postsynaptic 13A | https://neuromancer-seung-import.appspot.com/?json_url=https://raw.githubusercontent.com/sagrawal/Lee_2024/main/jsons/DS_13A.json |
| Postsynaptic 13B | https://neuromancer-seung-import.appspot.com/?json_url=https://raw.githubusercontent.com/sagrawal/Lee_2024/main/jsons/DS_13B.json |
| Postsynaptic 14A | https://neuromancer-seung-import.appspot.com/?json_url=https://raw.githubusercontent.com/sagrawal/Lee_2024/main/jsons/DS_14A.json |
| Postsynaptic 18B | https://neuromancer-seung-import.appspot.com/?json_url=https://raw.githubusercontent.com/sagrawal/Lee_2024/main/jsons/DS_18B.json |
| Postsynaptic 19A | https://neuromancer-seung-import.appspot.com/?json_url=https://raw.githubusercontent.com/sagrawal/Lee_2024/main/jsons/DS_19A.json |
| Postsynaptic 19B | https://neuromancer-seung-import.appspot.com/?json_url=https://raw.githubusercontent.com/sagrawal/Lee_2024/main/jsons/DS_19B.json |
| Postsynaptic 20A | https://neuromancer-seung-import.appspot.com/?json_url=https://raw.githubusercontent.com/sagrawal/Lee_2024/main/jsons/DS_20A.json |
| Postsynaptic 21A | https://neuromancer-seung-import.appspot.com/?json_url=https://raw.githubusercontent.com/sagrawal/Lee_2024/main/jsons/DS_21A.json |
| Postsynaptic 22A | https://neuromancer-seung-import.appspot.com/?json_url=https://raw.githubusercontent.com/sagrawal/Lee_2024/main/jsons/DS_22A.json |
| Postsynaptic 23B | https://neuromancer-seung-import.appspot.com/?json_url=https://raw.githubusercontent.com/sagrawal/Lee_2024/main/jsons/DS_23B.json |

| | |
|---|---|
| Postsynaptic motor neurons | https://neuromancer-seung-import.appspot.com/?json_url=https://raw.githubusercontent.com/sagrawal/Lee_2024/main/jsons/MN.json |
| Postsynaptic unknown | https://neuromancer-seung-import.appspot.com/?json_url=https://raw.githubusercontent.com/sagrawal/Lee_2024/main/jsons/unknown.json |

**Supplemental Table 3.** Neuroglancer links to ascending club, 8B, and 10B neurons in Flywire

| | |
|---|---|
| Ascending club neurons | https://ngl.cave-explorer.org/ - !middleauth+https://global.daf-apis.com/nglstate/api/v1/6680057844072448 |
| Ascending 8B/10B neurons | https://ngl.cave-explorer.org/#!middleauth+https://global.daf-apis.com/nglstate/api/v1/4958033236983808 |
